# Deep-learning based fully automatic segmentation of the globus pallidus interna and externa using ultra-high 7 Tesla MRI

**DOI:** 10.1101/2020.10.15.341578

**Authors:** Oren Solomon, Tara Palnitkar, Remi Patriat, Henry Braun, Joshua Aman, Michael C. Park, Jerry Vitek, Guillermo Sapiro, Noam Harel

**Affiliations:** Center for Magnetic Resonance Research, Department of Radiology, University of Minnesota, 2021 6^th^ Street S.E. Minneapolis, MN, 55455, USA; Department of Neurology, University of Minnesota, 516 Delaware St. S.E., 12-100 Phillips Wangensteen Building, Minneapolis, MN, 55455, USA; Department of Neurosurgery, University of Minnesota, MMC 96, Room D-429, 420 Delaware St. S.E, Minneapolis, MN, 55455, USA; Department of Electrical and Computer Engineering, Department of Biomedical Engineering, Department of Computer Science, Department of Mathematics, Duke University, Durham, NC, USA

**Keywords:** Deep-Learning, Convolutional Neural Network, Segmentation, Globus Pallidus, Deep Brain Stimulation, Parkinson’s Disease, Patient-Specific, 7 Tesla Magnetic Resonance Imaging

## Abstract

Deep brain stimulation (DBS) surgery has been shown to dramatically improve the quality of life for patients with various motor dysfunctions, such as those afflicted with Parkinson’s disease (PD), dystonia, and essential tremor (ET), by relieving motor symptoms associated with such pathologies. The success of DBS procedures is directly related to the proper placement of the electrodes, which requires the ability to accurately detect and identify relevant target structures within the subcortical basal ganglia region. In particular, accurate and reliable segmentation of the globus pallidus (GP) interna is of great interest for DBS surgery for PD and dystonia. In this work, we present a deep-learning based neural network, which we term GP-net, for the automatic segmentation of both the external and internal segments of the globus pallidus. High resolution 7 Tesla images from 101 subjects were used in this study; GP-net is trained on a cohort of 58 subjects, containing patients with movement disorders as well as healthy control subjects. GP-net performs 3D inference in a patient-specific manner, alleviating the need for atlas-based segmentation. GP-net was extensively validated, both quantitatively and qualitatively over 43 test subjects including patients with movement disorders and healthy control and is shown to consistently produce improved segmentation results compared with state-of-the-art atlas-based segmentations. We also demonstrate a post-operative lead location assessment with respect to a segmented globus pallidus obtained by GP-net.

## 1. Introduction

In the past several decades, deep brain stimulation (DBS) therapy has shown clear clinical efficacy in the mediation of symptomatic motoric behavior associated with Parkinson’s disease (PD), essential tremor (ET), dystonia, and other conditions (Benabid et al., 1987; Deuschl et al., 2006; Hariz et al., 2008; Mueller et al., 2008; Obeso et al., 2001; Volkmann et al., 2012). One of the most prominent DBS targets for PD and dystonia is the globus pallidus (GP) (Patriat et al., 2018). The GP is divided into two compartments, the internal segment (GPi) and the external segment (GPe), of which the former is typically the actual target for electrode placement. The GPe and GPi are separated by a thin layer, called the internal medullary lamina (Lozano & Hutchinson, 2002; Patriat et al., 2018). Several past studies have reported that lesions applied to the GPi led to improvement in motor function (Baron et al., 1996; Obeso et al., 2001; Vitek et al., 2003). The application of lesions (or pallidotomy) is associated with non-reversible risks of applying the lesion outside of the intended target. DBS surgery, on the other hand, has risen as an alternative with similar benefits, whose application can be reversed or even stopped if erroneously applied to the wrong region (Benabid et al., 1987; Obeso et al., 2001). Recent studies have shown that accurate placement of the DBS electrode within the sensorimotor region of the target (e.g. subthalamic nucleus (STN) or GPi) is directly correlated with the success of the DBS procedure and reduction of adverse effects (Ellis et al., 2008; Marks et al., 2009; Paek et al., 2013; Patel et al., 2015; Richardson et al., 2009; Rolston et al., 2016; Welter et al., 2014). Correct anatomical target identification is characterized not only by the target’s center of mass, but also by its boundaries (Kim et al., 2019). Thus, precise identification of both GPe and GPi and their lamina boundary is of great importance.

A fully automated segmentation process of the GP (both the internal and external segments) has several clear advantages, among which are accurate and fast inference. From a clinical point of view, an automated process has the potential to streamline clinical workflow and increase patient throughput, both in pre-operative surgery planning and post-operative assessment of the DBS lead location with respect to the target. Such a process can also eliminate human bias associated with the segmentation process, and provide more accurate and consistent segmentation results.

Since some anatomical structures are not easily identified or visualized on standard clinical images (e.g. 1.5 Tesla (T) or 3 T MRI scanners), a common approach to localize brain structures, and in particular those located in the basal ganglia, is to rely on an atlas (Ewert et al., 2018; Horn et al., 2019; Horn & Kühn, 2015)^i^. An atlas provides an average location of brain structures, often based on multiple inputs, such as different MRI scans and histology, merged from numerous subjects. For example, the authors of (Chakravarty et al., 2006) combined a histological atlas with a 3 T based multimodal subcortical atlas built from MRIs of PD patients (Xiao et al., 2017). In addition, they also combined high resolution multimodal MRIs and structural connectivity data as well (Ewert et al., 2018). Atlases can be deterministic (Xiao et al., 2017), that is, each pixel corresponds to a single brain structure, or probabilistic (Ewert et al., 2018), where each pixel is associated with a vector of probabilities which indicated how likely that pixel is associated with different brain structures. For example, (Horn, Kühn, et al., 2017) took a probabilistic approach to map DBS electrode locations onto the Montreal Neuorological Institute (MNI) space.

Although atlases, which are typically defined in a normalized space, have shown great importance in retrospective population studies (Horn, Kühn, et al., 2017; Horn, Neumann, et al., 2017; Horn, Reich, et al., 2017; Kim et al., 2019), several recent studies have shown that variablity exists of the size and shape of deep brain structures between different subjects (Abosch et al., 2010; Duchin et al., 2018; Lenglet et al., 2012; Patriat et al., 2018). This inherent inter-patient variability is not fully captured by atlas-based segmentations, as they provide a single, averaged shape of brain structures, be it deterministic or probabilistic. Accounting for inter-patient variablity is typically done by registering the atlas to specific patient anatomy (i.e. via an MRI scan), although this approach cannot often account for the discrepancy between the template and the individual patient brain, and is also affected by registration errors (Dadar et al., 2018; Ewert et al., 2018; Kim et al., 2019). Several other approaches have been introduced in recent years. However, these approaches segment the GP as an entire structure and do not provide distinction between the GPe and the GPi (Manjón & Coupé, 2016; Visser et al., 2016). These drawbacks motivate the development of a truly patientspecific segmentation technique for GPe/GPi structures.

In recent years, the fields of image analysis and computer vision have undergone a monumental and profound change with the introduction of deep-learning (DL), and in particular deep fully convolutional neural networks (CNNs) (Krizhevsky et al., 2012; Lecun et al., 2015). These types of networks consist of many aggregated layers of convolution filters of various sizes and non-linear elementwise activation functions, such as the rectified linear unit (ReLU). In many computer vision tasks (e.g. segmentation/classification), CNNs are trained end-to-end in a fully supervised manner, over pairs of input images and (often manual) delineations or labels. Such trained CNNs have shown state-of-the-art performance in many tasks, such as semantic segmentation (Long et al., 2015), image classification (Krizhevsky et al., 2012; Zhong et al., 2015), and image registration (Balakrishnan et al., 2018; Dalca et al., 2018) to name a few. As opposed to iterative algorithms, CNNs perform inference in a single forward step without any need for time-consuming iterations. Moreover, since CNNs are composed from convolution operations and elementwise non-linearities, they are implemented very efficiently, which leads to fast execution times during inference.

In this work we present *GP-net*^ii^, a fully convolutional deep neural network for the efficient and accurate 3D segmentation of both the GPe and GPi. GP-net is based on a variant of one of the most prominent deep networks architecture, the U-net (Ronneberger et al., 2015), which exploits skip connections in order to prevent loss of contextual information at multiple image scales. In particular, we exploit the attention gated (AG) U-net proposed in (Schlemper et al., 2019), which has previously shown improved segmentation performance in medical imaging. Attention mechanisms are able to automatically learn to focus, or direct attention, without additional supervision. This ability allows AGs to highlight salient features during inference time in the input images or intermediate feature maps, and have been applied successfully in numerous machine learning disciplines, such as natural language processing and machine vision (Bahdanau et al., 2015; Wang & Shen, 2018). For example, in (Schlemper et al., 2019) it was shown that the added AGs improve model sensitivity and accuracy in medical computerized tomography (CT) and ultrasound segmentation, by suppressing irrelevant feature activations in irrelevant areas.

In addition, we augment the attention gated U-net with the recently introduced deformable convolutions (Dai et al., 2017; Pominova et al., 2019), by replacing some of the intermediate 3D convolution layers in the network with 3D deformable convolutions. Classical convolution filters rely on convolving the learned kernel with input which lies on a regular Cartesian grid. In (Dai et al., 2017) it was shown that by learning the grid samples, instead of using a fixed grid, improved performance in vision-based tasks such as classification and segmentation can be achieved. In that sense, deformable convolutions can be thought of as another form of an attention mechanism.

Recent advances in ultra-high MRI machines and acquisition protocols have allowed 7 T imaging methods to directly visualize and identify small subcortical deep brain structures. Structures such as the STN or the lamina border between the GPe and GPi cannot be clearly visualized in lower field MRI machines, but can be more clearly identified with the use of 7 T due to improved contrast and resolution (Abosch et al., 2010; Patriat et al., 2018). Ultra-high field MR have already been used for deep brain structure identification; for example (Duchin et al., 2018) have shown that direct visualization of the STN is possible, and (Kim et al., 2019) have shown that by relying on 7 T manual delineations and machine learning techniques, direct segmentation of the STN on 3 T images is possible, with 7 T accuracy and precision. Atlas-based approaches have also exploited the benefits of 7 T imaging and superior contrast to constrcut state-of-the-art altases. One such example is the work of Keuken et. al. (Keuken et al., 2014) which constructed an atlas which relies on multi-contrast 7 T acquisitions to analyze the anatomincal variablity of subcortical structures.

In this work, we rely on acquired T2 volumes from ultra-high 7 T MRI machine, specifically tailored to visualize the basal ganglia region. The training cohort consists of movement disorder patients, as well as healthy control subjects, such that for each subject manual delineations of both left and right GPe and GPi are obtained by experts to train the network end-to-end. An overall illustration of the proposed process is given in Fig. 1. In the next sections we outline the mechanism behind GP-net, provide extensive experimental validation and finish with a discussion and concluding remarks.

**Figure 1.**
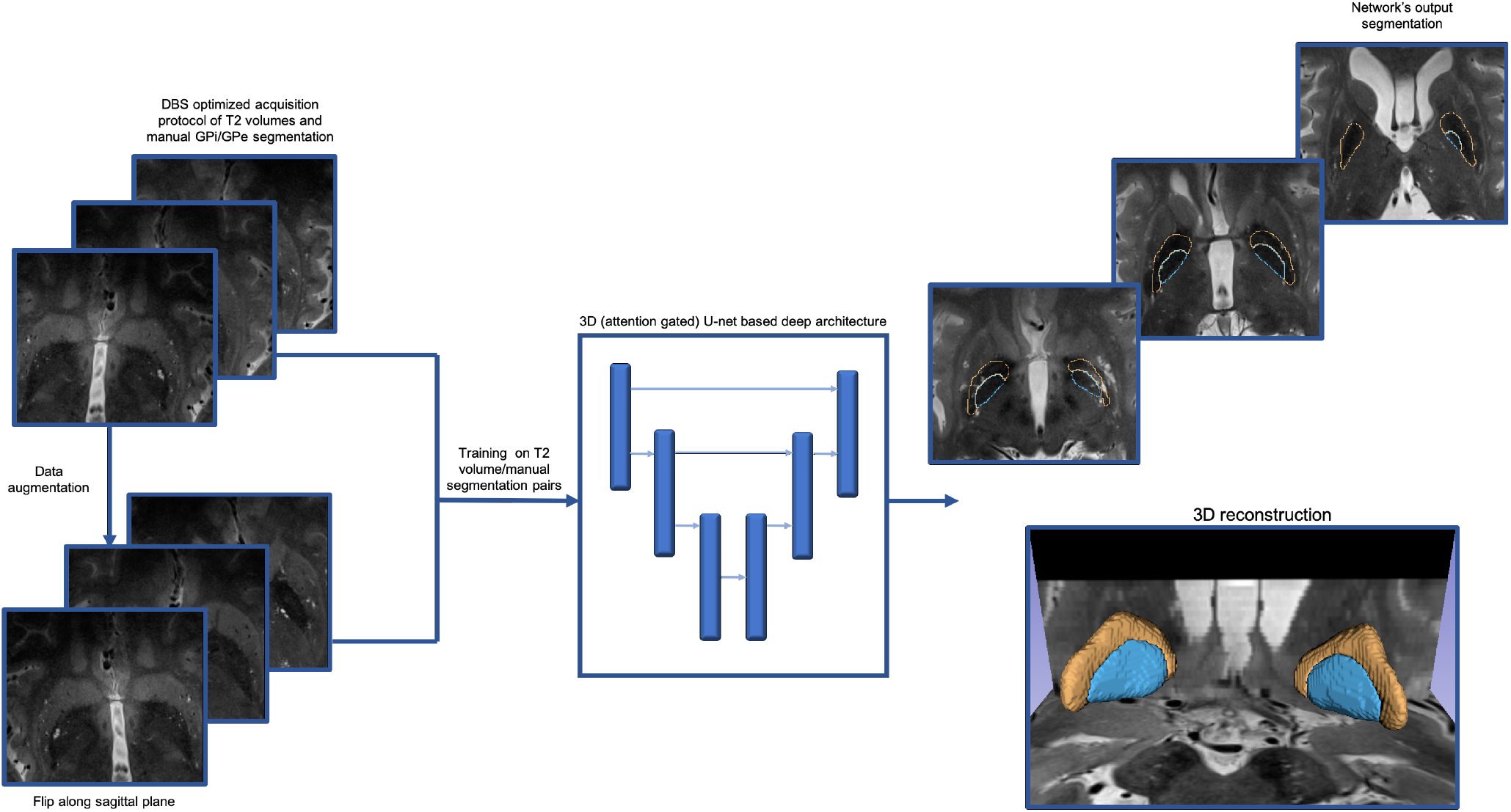
Proposed GP-net for the automatic and patient-specific segmentation of the GPe and GPi from acquired 7 T T2 volumes. GP-net is an attention gated U-net, trained end-to-end from pairs of 7 T T2 scans and manual 3D delineations of both GPe and GPi. To increase training size, each training pair is augmented by flipping the images along its sagittal plane. Resulting network’s output are the automatically segmented GPe (orange) and GPi (blue) 3D volumes.

## 2. Methods

### 2.1. Overview

GP-net is trained end-to-end in a fully supervised manner. Manual 3D delineations performed by domain experts, for both the GPe and GPi are extracted per each patient’s 7 T T2 scan. Thus, the network is fed with pairs of 7 T T2 volumes and corresponding 3D manual segmentations in the training procedure.

To evaluate the performance of GP-net, as well as the quality of its patient-specific segmentations, the resulting automatic delineations are compared against manual GP segmentations, performed on 7 T T2 scans by experts, from a test set, which was excluded from the training phase of the network. The following metrics against the manual delineations are compared: dice score, center of mass Euclidean distance, volume and mean surface differences. In addition, we provide extensive quantitative comparison between GP-net and four state-of-the-art atlases. GP-net shows significant improvement over existing atlas-based techniques. A test-retest experiment was performed to assess the consistency of GP-net in segmenting the same structure of the same patient over several different scans acquired over the course of days. Finally, to evaluate the clinical utility of the proposed method we compared the DBS lead location post-surgery as determined based on manual segmentation and the automatic GP-net segmentations.

### 2.2. Scanning protocol

Patients were scanned on a 7 T MRI scanner (Magnetom 7 T Siemens, Erlangen, Germany) using our previous published protocols (Abosch et al., 2010; Duchin et al., 2018). The scanner was equipped with SC72 gradients capable of 70 mT/m and a 200 T/m/s slew rate using a 32-element head array coil (Nova Medical, Inc., Burlington, MA, USA). On the day of scanning, the patients were instructed to take their usual medication in order to optimize patient comfort and minimize motion. Whenever patient head size enabled enough space in the coil, dielectric pads were utilized in order to enhance signal in the temporal regions (Teeuwisse et al., 2012). The scan protocol consists of: T1-weighted whole brain scan (0.6 mm^3^ isotropic) and T2-weighted axial slab covering from the top of the thalamus to the bottom of the substantia nigra with 0.39 × 0.39 × 1 mm^3^ resolution. The T1 weighted scan was used only for atlas-based registration when comparing GP-net with different atlases and was not used in the network’s training, inference or validation phases (details are given in subsection 2.6 and the Results section).

### 2.3. Database and pre-processing

A cohort of 101 subjects, including 24 healthy controls and 77 movement disorder (PD and ET) patients participated in this study. All subjects were scanned on the 7 T scanner; patients were scanned prior to their DBS surgeries. Even though ET patients do not typically undergo GPi-DBS, we also chose to include their imaging data in our dataset. Out of the 101 participatens, images from 58 participants were used for training and 43 for testing; demographic details are given in Table 1. In addition, one subject was scanned three times on two days with the same scanning protocol to assess the network’s stability. For each subject in the cohort, manual delineation of both the left and right GPe and GPi were obtained by three independent experts from the scanned T2 volumes.

**Table 1.** **A)** Subjects demographics. ET = Essential Tremor, PD = Parkinson’s Disease. Age was determined at the day of the scan. **B)** Subjects mean (av) age ± standard deviations (std), divided into training and testing.

|  | *Ages 20-40* |  | *Ages 41-60* |  | *Ages 61-80* |  |
| --- | --- | --- | --- | --- | --- | --- |
|  | **Training** | **Testing** | **Training** | **Testing** | **Training** | **Testing** |
| *CONTROL* | 9 | 0 | 2 | 7 | 1 | 5 |
| *ET* | 0 | 0 | 7 | 0 | 8 | 8 |
| *PD* | 0 | 0 | 11 | 8 | 20 | 15 |

| *CONTROL* |  | *ET* |  | *PD* |  |
| --- | --- | --- | --- | --- | --- |
| **Training** | **Testing** | **Training** | **Testing** | **Training** | **Testing** |
| 34.83 ± 15.68 | 59 ± 8.95 | 63.2 ± 9.45 | 68.75 ± 4.33 | 63.93 ± 8.11 | 64.08 ± 9.71 |

To increase the number of training pairs (T2 image and manual delineations), each training pair was mirrored along the sagittal plane (Fig. 1). Thus, the number of training pairs was doubled and provided more training examples of spatially translated GP structures. The T2 volumes and manual segmentations were resampled to an isotropic grid of 0.39 × 0.39 × 0.39 mm^3^ with a nearest neighbor interpolation kernel prior to training and inference. All of the quantitative analysis is performed on the resampled grid. This study was approved by the Institutional Review Board at the University of Minnesota and all participants gave their informed consent.

### 2.4. Network architecture

As previously mentioned, GP-net is a fully convolutional 3D deep neural network. Its base architecture consists of an attention gated U-net, in which some of the inner 3D convolution layers were replaced by 3D deformable convolutions. The basic U-net architecture is composed of two paths, encoder and decoder paths, which consist of aggregated layers of convolutions, max-pooling, and ReLU activations. Each layer shrinks its input size by a factor of 2. Thus, at the end of the encoder stage, the feature dimensions are shrunk by a factor of 2^*n*^, where *n* is the number of encoder stages (assuming isotropic max-pooling in all input dimensions). The output of this stage is then fed into the decoder, which consists of the same number of *n* layers, each layer is built with deconvolution layers and upsampling by a factor of 2. The final output of the network has the same dimensions as the input. In this work, *n* = 4. Each encoder stage is composed of two consecutive blocks of 3D convolution (same parameters in both blocks), 3D batch normalization, ReLU activation followed by a max pooling layer of size 2 × 2 × 2 pixels. Deformable convolution layers consist of a 3D standard convolution kernel to estimate the offsets, a deformable convolution layer, followed by another standard 3D convolution (all convolutions of same kernel size and padding). Detailed parameters of the convolution layers are given in Table 2.

**Table 2.** GP-net convolution layers parameters. Kernel sizes are isotropic. All numbers are given in pixels. There is no dilation for the deformable convolution kernels.

| *Stage* | *Block level* | *Type* | *Kernel size* | *Padding size* | *Dilation* |
| --- | --- | --- | --- | --- | --- |
| *Encoder* | 1 | Regular 3D Convolution | 11 | 15 | 3 |
|  | 2 | Regular 3D Convolution | 9 | 8 | 2 |
|  | 3 | Deformable 3D Convolution | 5 | 2 | - |
|  | 4 | Deformable 3D Convolution | 3 | 1 | - |
| *Central* | 5 | Deformable 3D Convolution | 3 | 1 | - |
| *Decoder* | 4 | Deformable 3D Convolution | 3 | 1 | - |
|  | 3 | Deformable 3D Convolution | 3 | 1 | - |
|  | 2 | Regular 3D Convolution | 5 | 2 | 1 |
|  | 1 | Regular 3D Convolution | 7 | 3 | 1 |
| *Final* |  | Regular 3D Convolution | 1 | 0 | 1 |

In addition, as illustrated in Fig. 1, each layer in the encoder stage is directly connected to its corresponding decoder stage, traditionally using skip connections. Skip connections allow more efficient gradient flow through the network during the training stage and prevent loss of contextual information at multiple image scales. Following (Schlemper et al., 2019), in this work the encoder stages are connected to the corresponding decoder stages via attention gates. A detailed description of the attention gate architecture, as well as the overall network architecture is given in (Schlemper et al., 2019). GP-net has three attention gates, for the second, third, and fourth layers of the decoder stage. Each block in the decoder stage is composed of a 3D convolution kernel, followed by an upsampling operator by a factor of 2 × 2 × 2. At each decoder level, the resulting attention signal and corresponding upsampled feature map from the previous lower decoder stage are concatenated and convolved with a 3D convolution filter of kernel size 1, no padding and no dilation. Deformable convolutions in the decoder stage are structured the same way as deformable convolutions in the encoder stage. Between the encode and decoder there is another stage (central stage) which is used as the gating signal for the fourth AG.

### 2.5. Training loss function

GP-net is trained end-to-end using pairs of T2 volumes and corresponding manual delineations. The training loss function used in this work is a combination of several loss functions, detailed below.

Tversky loss (Salehi et al., 2017): In the case of binary segmentation (e.g. network’s output and manual delineation), the Tversky index between two groups *A* and *B* is written as

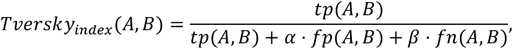

where *tp* stands for true positives (correctly classified voxels), *fp* stands for false positives (wrongly classified voxels), *fn* stands for false negatives (wrongly missclassified voxels), and *α* and *β* are corresponding weights. The Tversky loss is taken as 1 - *Tversky_index_*, as we wish to minimize the loss function through gradient descent. The Tversky loss was reintroduced and utilized in the context of deep-learning based segmentation (Kim et al., 2020; Salehi et al., 2017) as an efficient tool for handling imbalanced class labels, since the parameters *α* and *β* control the relative weights between false positives and false negatives. In this work we choose these parameters according to Table 3.

**Table 3.**
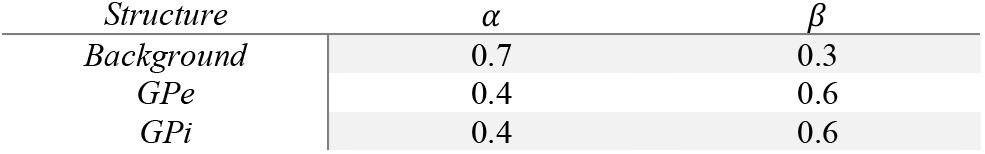
Tversky loss parameters.

The Tversky index can be considered as a generalization of the dice index (Dice, 1945), and indeed by taking *α* = *β* = 0.5 the Tversky index coincides with the dice index. Additionally and similarly to (Kim et al., 2019), to minimize possible overlap between two different classes in the segmentation, we penalize (minimize) the dice score between each prediction to each other label. This term is weighed by a factor of 0.01 relative to the Tversky loss.

Hausdorff distance (Karimi & Salcudean, 2020): The Hausdorff distance measures the largest (in our case Euclidean) distance between two given contours. Thus, minimizing the Hausdorff distance can be though of as minimizing the worst case, or largest outlier distance between the network’s segmentation and the manual segmentation, which is indicative of the largest segmentation error. Given two point sets *X* and *Y,* the one sided Huasdorff distance is defined as (Karimi & Salcudean, 2020; Rockafellar & Wets, 2009)

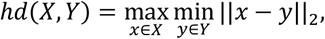

and the bidirectional Hausdorff distance is given by

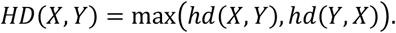

Since the Hausdorff metric is highly non-differentiable, we require some differentiable proxy in order to use back-propagation. We rely on the proposed estimator given in Eq. (8) of (Karimi & Salcudean, 2020). This estimator is a smooth approximation of the Hausdorff distance, which allows back-propagation using gradient descent. In practice, we found it most efficient to minimize the Hausdorff distance for the entire GP (left and right sides together). Initial weight relative to the Tversky loss is 0.00001 and increases by a factor of 5 every 50 epochs.

We train the network of 94 epochs using stochastic gradient descent with learning rate of 0.0001 and momentum factor of 0.9. Batch size is 1. GP-net was implemented in Python 3.6 with PyTorch 1.4 and trained on a single Nvidia V100 GPU with 32 GBs of memory.

### 2.6. Validation

To evaluate GP-net, we performed an extensive quantitative analysis of its performance over 43 subjects. These subjects were not included in the training cohort, and were only used for inference and evaluation of the network’s performance.

GP-net is compared to the manual segmentation performed by domain experts and against four publicly available state-of-the-art atlases, to quantify its performance and validate its reliablity for patient-specific GPe/GPi segmentation: (1) DBS intrinsic template atlas (DISTAL, referred to as Ewert 2017) (Ewert et al., 2018); (2) California Institute of Technology (CIT) probabilistic high-resolution in vivo atlas of the human amygdala, also known as the CIT168 reinforcement learning atlas (referred to as Pauli 2020) (Pauli et al., 2018); (3) population-averaged atlas that was made with 3 T MRI of 25 PD patients (referred to as Xiao 2017) (Xiao et al., 2017); and (4) atlas of the basal ganglia and thalamus (referred to as He 2020) (He et al., 2020). All atlases were taken from the lead-DBS software package (Horn et al., 2019; Horn & Kühn, 2015), in which they are registered to the MNI ICMB2009b template [5].

Since GP-net is patient-specific and operates on the patient’s T2 (isotropically resampled) volumetric scan, to perform a fair comparison, we first register the atlases to the same T2 space. This registration process starts with a registration of the T1 MNI ICBM2009b (3 T) template to a 0.39 mm isotropic MNI template (3 T) using the Advanced Normalization Toolbox (ANTs) (Avants et al., 2011), via the command antsApplyTransforms and the LanczosWindowedSinc interpolation kernel. Next, the registered template is registered to the patient-specific (0.39 mm isotropically resampled) T1 scan (7 T). This is done via a combination of FLIRT and FNIRT modules from the FSL toolbox (Jenkinson et al., 2012) (implemented via HCP pipelines), using the ApplyWarp command and a spline interpolation kernel. The final registration stage includes registering the patient’s T1 scan into his/her (isotropically resampled) T2 scan (7 T) using ANTs with a B-spline interpolation kernel and linear registration. The same transformations are applied to the atlases with a nearest-neighbor interpolation kernel. All registrations were verified visually.

We note that some of the atlases, such as the DISTAL (Ewert 2017 (Ewert et al., 2018)) and CIT168 (Pauli 2017 (Pauli et al., 2018)) are probabilistic, while GP-net provides a deterministic segmentation map. To make a fair comparison, we have thresholded these atlases with a value of 0.001, meaning that for each class (GPe/GPi), voxels with values below 0.1% probability are zeroed out, while voxels with higher probability are given a value of 1 (GPe) or 2 (GPi). This threshold value was validated visually for each atlas and compared with the T2 ICBM2009b template. We also tried different values (such as 0.01), which yielded similar quantitative results.

On a final note, in some segmentation cases, we observe that the output of GP-net might contain small segmented regions (“islands”) which are clearly not related to either the GPe or GPi. These regions are easily removed automatically with a small post-processing step which removes all segmentation regions except for the largest four (left and right GPe and GPi) through the use of the connected components algorithm (Rosenfeld & Pfaltz, 1966).

### 2.7. Metrics and statistics

We use the following metrics to compare between the performance of GP-net and the different atlases: (1) Dice score, (2) Center of Mass distance (CoM) (mm), (3) Mean (Euclidean) Surface Distance (MSD) (mm), (4) volume estimation (mm^3^), and (5) precision vs. recall rates. Dice, CoM and MSD are also calculated against the manual GPe and GPi segmentations. CoM distance describes how well the segmented structure is being localized in space (i.e. inside the brain), while the dice, MSD and volume meaurements dscribe how well the shape of the structure is being captured by the different segmentation techniques.

A one-way analysis of variance (ANOVA) was calculated for each metric, followed by a multiple comparison correction and post hoc tests with Tukey’s honest significant difference to determine statistical significance between the different methods.

The matrices (figs. 2 and 3) indicate the statistical significance between each method. Each cell in the matrices corresponds to the p-value that the method written in its corresponding row is statistically different from the method written in its corresponding column. A blue cell indicates p-value lower than 0.1%, a green cell indicates a p-value lower than 5% and a red cell indicates p-value higher than 5%.

**Figure 2.**
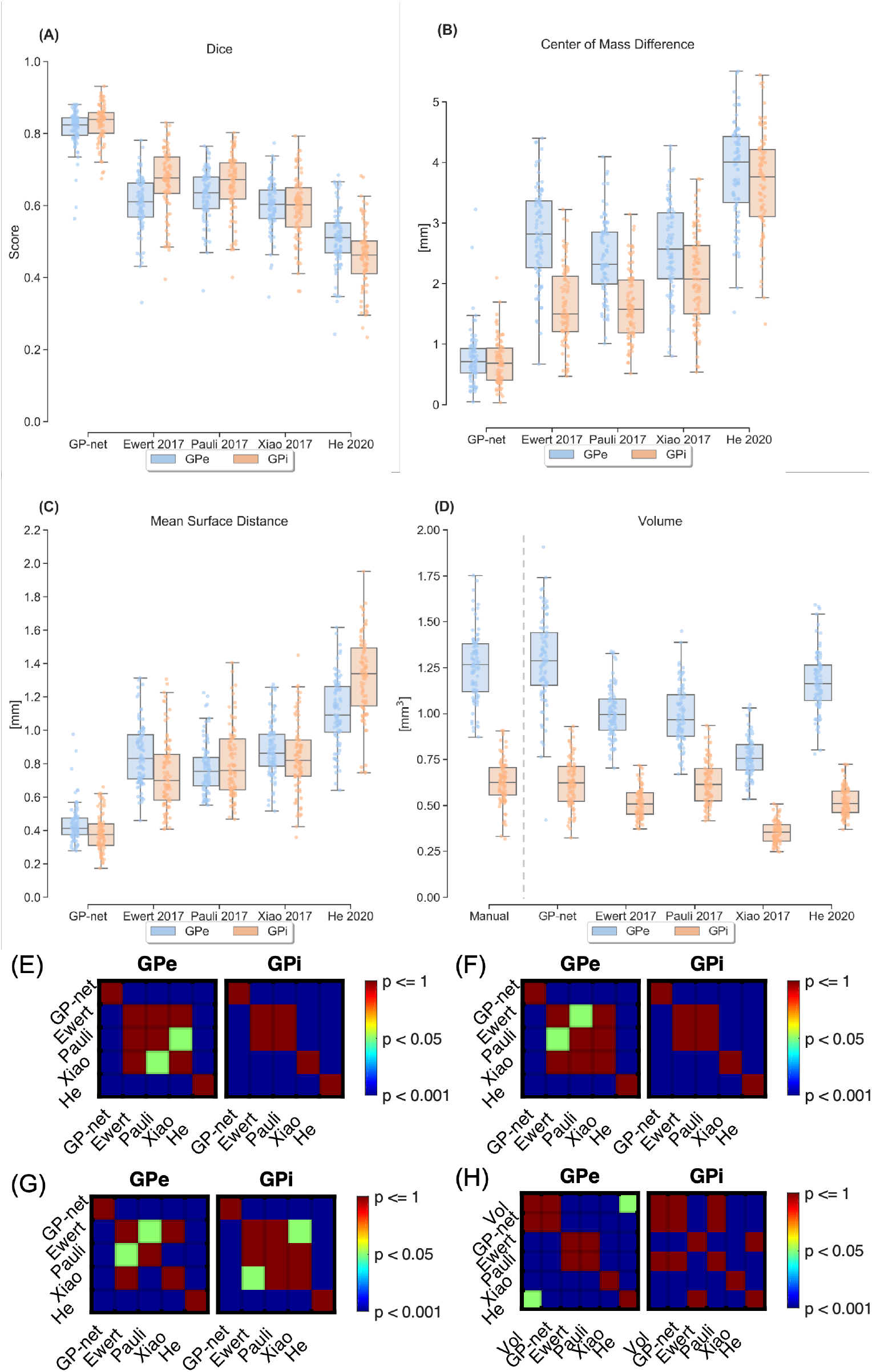
(A) Dice score comparison between GP-net and the different atlases for both GPe (blue) and GPi (orange) against manual segmentation. Values are compared against the manual delineations for both left and right sides of each patient and presented both as box plots and individual points. (B) Center of mass (CoM) distance relative to the manual delineations. (C) Mean surface distance (MSD) relative to the manual delineations. (D) Estimated volumes. Dashed vertical gray line in (D) separates between the manual volume estimates (left) and the different segmentation techniques estimates. (E) – (H) Statistical significance matrices between the different techniques for panels (A) – (D), respectively.

**Figure 3.**
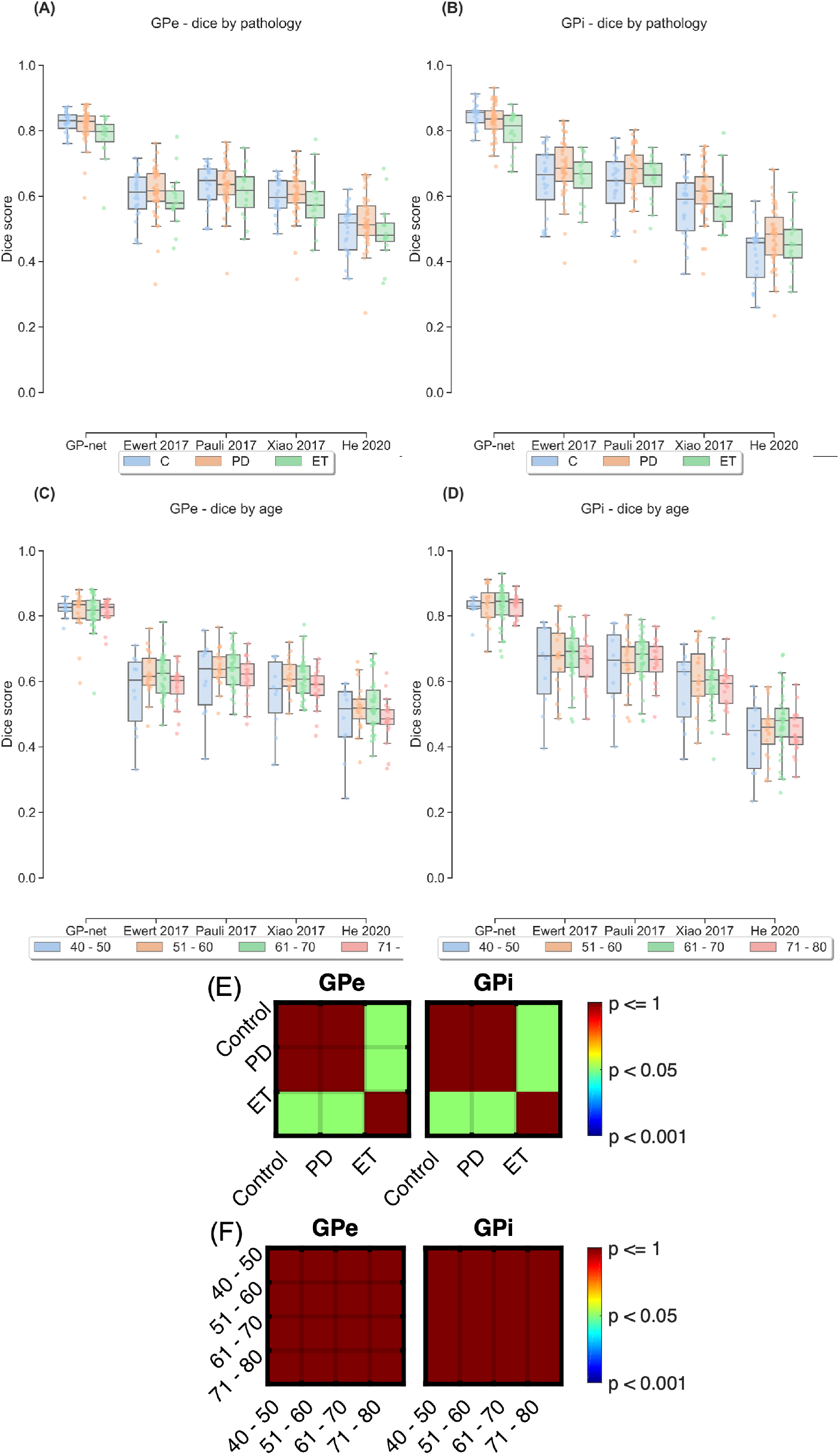
(A) and (B) dice scores divided by pathology type: healthy control in blue, PD in orange and ET in green for the GPe and GPi, respectively. (C) and (D) dice scores divided by age group (measured in years). (E) Statistical significance matrices for panels (A) and (B) between the healthy control, PD and ET cohorts for GP-net only. (F) Statistical significance matrices for panels (C) and (D) between the different age groups for GP-net only.

Additional metric which we utilize to compare between the different segmentations is the precision vs. recall rates of each method. Given a binary segmentation scenario, precision is defined as the ratio

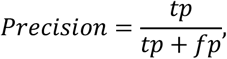

while recall (sensitivity) is defined as the ratio

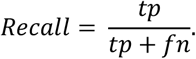

Precision measures a technique’s rate of correctly segmenting only GPe/GPi voxels as such, as opposed to classifying background voxels as either GPE or GPi (hence, to minimize false positives rate), while recall measures a technique’s rate of correctly capturing and segmenting GPe/GPi voxels as such, without misclassifying them as backgound (hence, to minimize false positives rate).

## 3. Results

In this section we present a detailed quantitative and qualitative analysis of the performance of GP-net, compared against state-of-the-art atlases and experts’ segmentation, as well as a stability test for GP-net.

### 3.1. Quantitative analysis

We start by presenting comparative results between GP-net and atlas-based segmentation, with respect to the manual delineations as can be seen in figs. 2 and 3, comparing the dice score, CoM (mm), MSD (mm), and volume (mm^3^). In all of these figures, the metrics are calculated per patient, per structure (GPe and GPi), and per side (left and right). Figure 2A shows dice scores for GP-net and the selected atlas-based segmentations, both for the GPe and GPi, individually. Panels (B), (C) and (D) show CoM difference, MSD difference, and volume estimates based on the different segmentation techniques, respectively. Matrices (E) – (H) indicate statistical significance (p-values) between each pair of the different methods for panels (A) – (D), respectively.

GP-net has notably superior performance relative to the atlas-based segmentations, with distinctly higher dice scores (for both GPe and GPi) and lower dispersity, as well as lower values for the average CoM difference, and MSDs. In both cases, p-value matrices show clear statistical significant difference between GP-net and the other atlas-based metrics. These results are further supported in Table 4 by lower mean values and standard deviations of the CoM and MSD measurements for GP-net compared with the atlas-based segmentations.

**Table 4.** Mean scores and standard deviations of dice scores, CoM, MSD and Volume (Vol) estimate comparisons between GP-net, atlas-based segmentation and 7 T manual segmentation (applicable only for volume estimate). *n* = 86.

|  |  | *7 T Manual* | ***GP-net*** | *Ewert 2017* | *Pauli 2017* | *Xiao 2017* | *He 2020* |
| --- | --- | --- | --- | --- | --- | --- | --- |
| *Dice* | GPe | - | **$0.81 \pm 0.05$** | $0.6 \pm 0.08$ | $0.62 \pm 0.07$ | $0.59 \pm 0.06$ | $0.51 \pm 0.08$ |
| | GPi | - | **$0.83 \pm 0.05$** | $0.66 \pm 0.08$ | $0.66 \pm 0.08$ | $0.6 \pm 0.08$ | $0.45 \pm 0.09$ |
| *CoM*<br>*[mm]* | GPe | - | **$0.76 \pm 0.45$** | $2.82 \pm 0.77$ | $2.42 \pm 0.64$ | $2.57 \pm 0.76$ | $3.84 \pm 0.79$ |
| | GPi | - | **$0.7 \pm 0.38$** | $1.63 \pm 0.64$ | $1.64 \pm 0.57$ | $2.1 \pm 0.75$ | $3.66 \pm 0.83$ |
| *MSD*<br>*[mm]* | GPe | - | **$0.44 \pm 0.11$** | $0.85 \pm 0.18$ | $0.77 \pm 0.15$ | $0.88 \pm 0.16$ | $1.11 \pm 0.2$ |
| | GPi | - | **$0.38 \pm 0.1$** | $0.74 \pm 0.21$ | $0.8 \pm 0.2$ | $0.84 \pm 0.21$ | $1.32 \pm 0.25$ |
| *Vol*<br>*[mm<sup>3</sup>]* | GPe | $1.25 \pm 0.19$ | **$1.29 \pm 0.24$** | $1.0 \pm 0.13$ | $1.0 \pm 0.16$ | $0.76 \pm 0.1$ | $1.17 \pm 0.16$ |
| | GPi | $0.63 \pm 0.12$ | **$0.62 \pm 0.13$** | $0.5 \pm 0.08$ | $0.62 \pm 0.12$ | $0.35 \pm 0.06$ | $0.52 \pm 0.07$ |

Figure 2(D) compares between GPe and GPi volume estimates. The two leftmost boxplots (left of the dashed gray vertical line) correspond to the volume estimates measured from the manual delineations, while the other columns correspond to the estimates measured by the different segmentation techniques. GP-net exhibits the closest similarity between the GPe and GPi volume distributions. Note that corresponding p-values between the manual estimates and GP-net’s estimates (both for the GPe and GPi) indicate no statistical differences. Most of the atlas-based estimations show statistically significant volume estimates errors, compared with the manual segmentations, and thus do not fully capture the entire volume distribution.

Figure 3 presents dice scores divided by disease and age groups. Panels (A) and (B) present the dice scores categorized according to condition (healthy control, PD, and ET), for the GPe and GPi, respectively, while panels (C) and (D) present the dice scores categorized according to four age groups. The two matrices in panel (E) indicate statistical significance (p-values), only for GP-net, between the different pathologies, for panels (A) and (B), respectively. Likewise, the two matrices in panel (F) indicate statistical significant difference (p-values), only for GP-net, between the different age groups for panels (C) and (D), respcetively. Panels (A) and (B) show that GP-net consistently achieves higher dice scores compared with the different atlas based segmentations, for healthy controls as well as PD and ET patients. Panels (C) and (D) demonstrate that GP-net’s performance is not affected by the large distribution of age amonst the scanned subjects (40 – 80 years). Atlas-based segmentations are not affected by age, however they exhibit lower dice scores than GP-net and higher distirubtion variance. GP-net is shown to consistently produce higher dice scores, without much variation among between and among the different age groups. Consistency of performance, as demonstrated for the proposed GP-net, is critical for DBS and for real deployment.

Lastly, we compare the precision vs. recall rates of each of the methods. Figure 4 presents the precision vs. recall rates of GP-net and atlas-based segmentations (both GPe and GPi). GP-net’s recall and precision rates are both approaching 1 (ideal case) and are higher than all other atlas-based segmentation rates. The average precision rates are 0.82±0.06, 0.71±0.08, 0.68±0.07, 0.8±0.08 and 0.51±0.08 for GP-net, Ewert 2017, Pauli 2017, Xiao 2017 and He 2020, respectively. The average recall rates are 0.82±0.08, 0.58±0.09, 0.61±0.09, 0.47±0.07 and 0.46±0.09 for GP-net, Ewert 2017, Pauli 2017, Xiao 2017 and He 2020, respectively. Moreover, the recall rates of GP-net are statistically significant from the recall rates of all other atlas-based segmentations (p-value < 0.001). Likewise, GP-net’s precision rates are statistically significant than all other atlas-based segmentations’ recall rates (p-value < 0.001), except from the precision rates of Xiao 2017.

**Figure 4.**
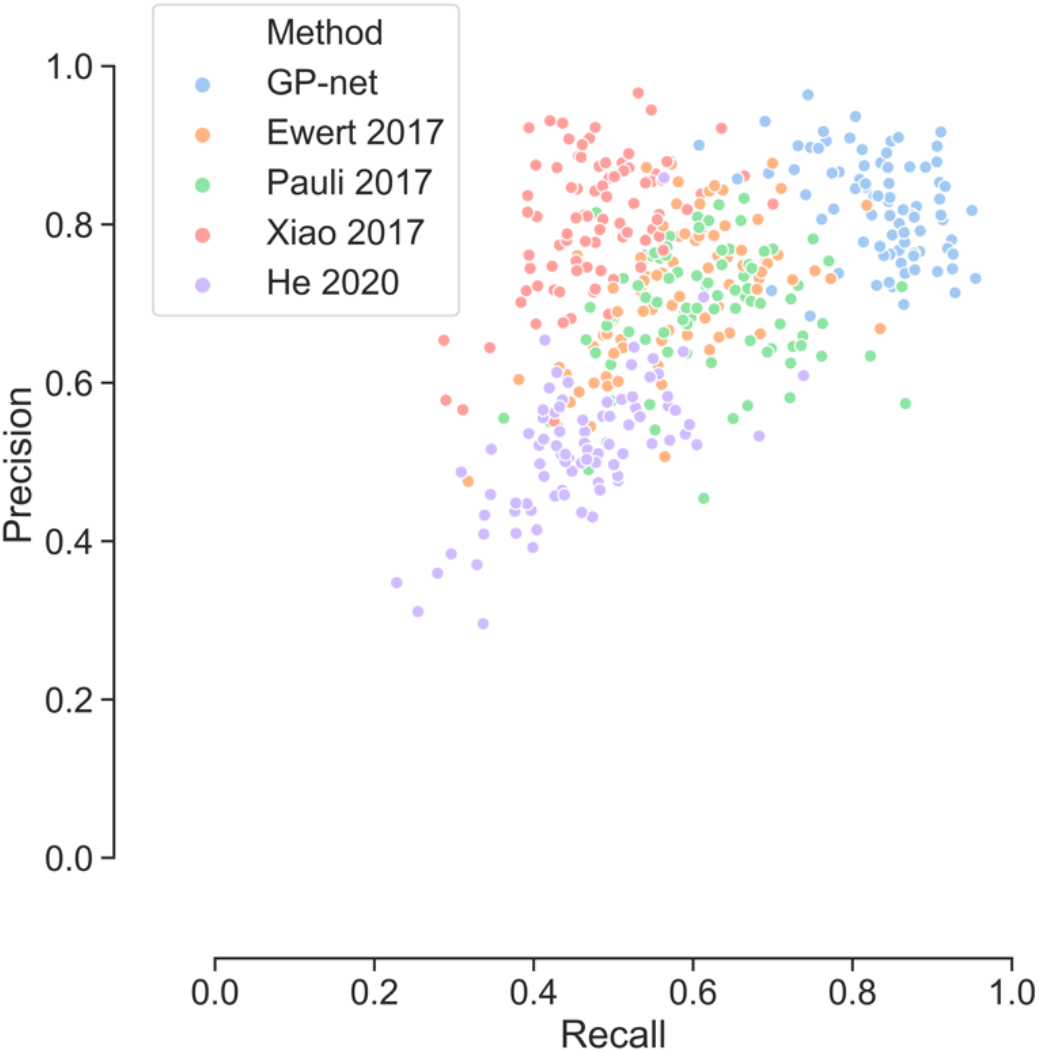
Precision vs. recall rates between the different methods (both GPe and GPi). Notice that GP-net does not exhibit a trade-off between the two rates and systematically achieves higher values than all of the other atlas-based segmentations.

### 3.2. Stability test

We further test the stability of the network to assess GP-net’s systematic behavior. We acquired three independent and repeated 7 T scans of a single healthy control subject over a period of two days, and performed inference using GP-net. Figure 5 presents the averaged (left and right hemisphers) dice, MSD, CoM and volume estimations for both GPe and GPi over the three scans.

**Figure 5.**
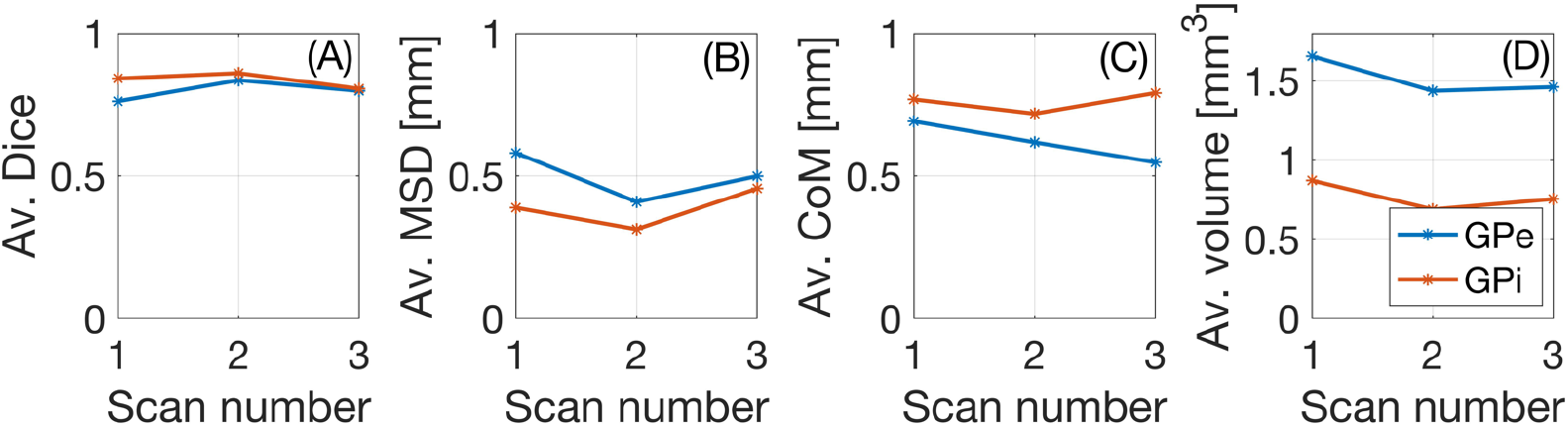
Test – retest scores. Av. indicates average over the left and right GPe (blue)/GPi (red). (A) Av. dice score, (B) av. MSD, (C) av. CoM and (D) av. estimated volume; GPe in blue and GPi in red.

The average deviations from the mean dice values in panel (A) are 3.09%/2.44% for the GPe and GPi respectively. The average deviations for the av. MSD (panel (B)) are 11.76%/12.56% for the GPe and GPi respectively, and for the av. CoM (panel (c)) are 7.78%/3.57% for the GPe and GPi respectively. Finally, the average deviations from the mean value for the volume estimation (panel (D)) are 6.06%/8.62% for the GPe and GPi respectively. These ranges are well inside the deviations reported in the literature for easier to compute brain characteristics, such as total volume. This analysis demonstrates GP-net’s ability to produce reliable and consistent segmentations, not only between different patients, as was previously shown, but also between different scans of the same subject over time.

### 3.3. Qualitative examples

Figure 6 illustrates segmentation results for a PD patient. The first column shows the 3D reconstructions of both left and right GPe (green for manual/ground truth (GT) and orange GP-net/atlas) and GPi (yellow for manual/GT and blue GP-net/atlas). Second and third columns show selected axial and coronal views (respectively) of a T2 slices, superimposed with the manual segmentation and GP-net/atlas segmentation, as well as the different metrics per each method. Judging visually, GP-net is able to produce the most accurate segmentation of both GPe and GPi, presenting excellent agreement with the manual delineation. Since GP-net performs inference directly on the isotropically resampled grid (0.39 mm^3^), its output is smooth, as can be seen in the 3D reconstruction, as opposed to the manual delineation (first row), which was segmented on the original 0.39 × 0.39 × 1 mm^3^ grid. To make a fair comparison, the atlas-based segmentations were registered from ICBM2009b to the isotropically resampled T2 grid through a 0.39 mm^3^ isotropic MNI template. However, even though we use the same grid spacing, these segmentation results seem to be more pixelated.

**Figure 6.**
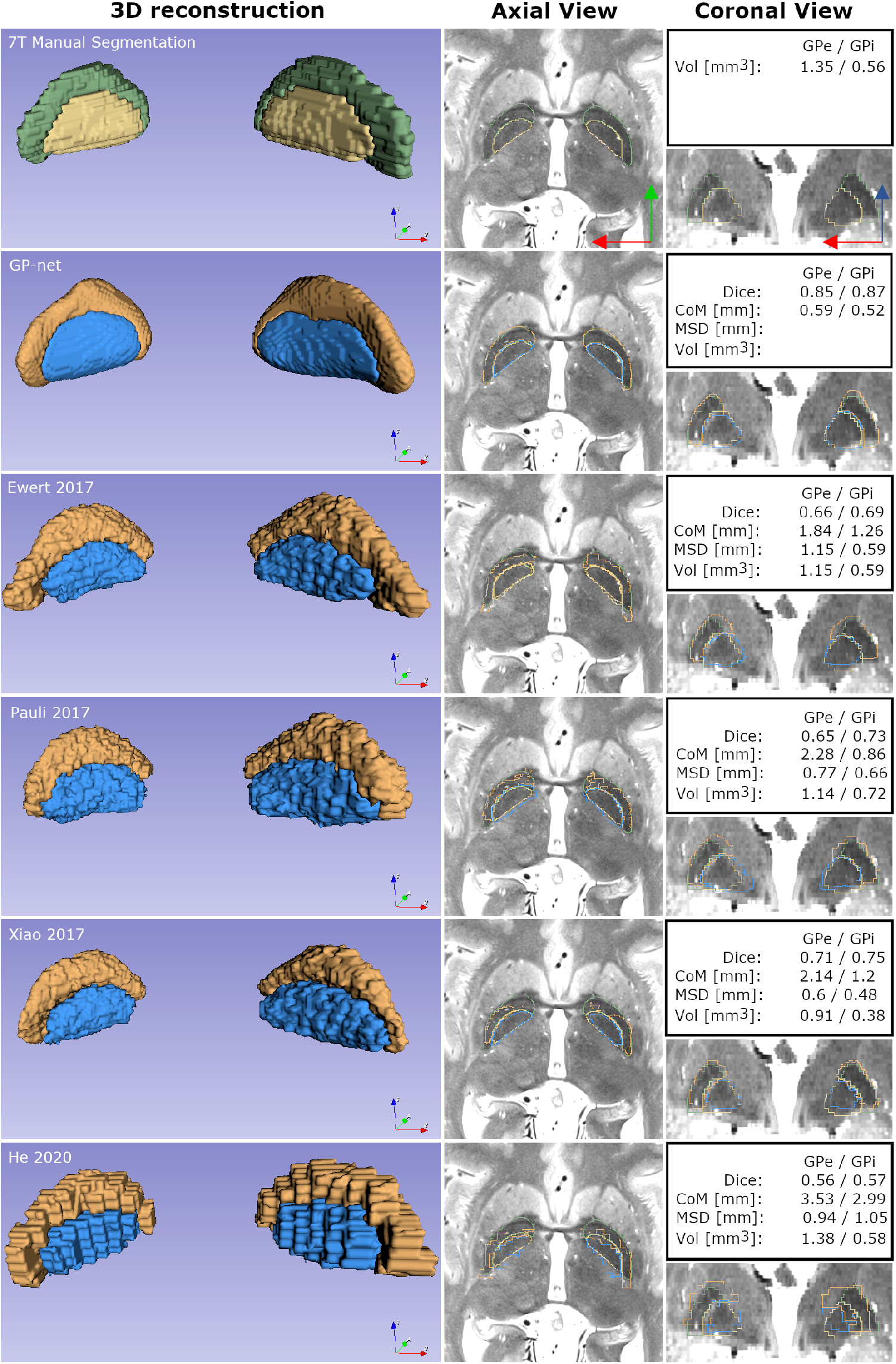
Segmentation and 3D reconstruction of the GP of a representitive PD patient. Leftmost column shows 3D reconstruction of both the GPe (green in the manual segmentation and orange in the different reconstructions) and GPi (yellow in the manual segmentation and blue in the different reconstructions). No smoothing was applied at any stage to the reconstructions or manual delineations. Middle column shows a selected T2 axial slice and superimposed outlines of manual delineation, the corresponding segmentations of GP-net (second row) and the different atlases (third – sixth rows). Rightmost column similarly illustrates a selected coronal view. Blue arrow points in the superior direction, green arrow points in the anterior direction and red arrow points in the right direction.

Figure 7 presents a clinical scenario, a retrospective anaysis of an actual GPi-targeted DBS case. Manual 7 T (panel (A)) and GP-net (panel (B)) 3D reconstractions of the GPe and GPi are shown. Electrode locations were extracted from a post-surgery CT acquired four weeks after the implant and registered with the MRI scan (Duchin et al., 2018). Excellent agreement of the electrode locations, with respect to the structures volumes and borders, as compared with the manual GPe and GPi delineations is observed between both panels. This image exemplifies the clinical potential of GP-net to produce high quality GPe and GPi segmentations from 7 T T2 scans for accurate and reliable visualization of image-based DBS targeting and post-surgery lead localization.

**Figure 7.**
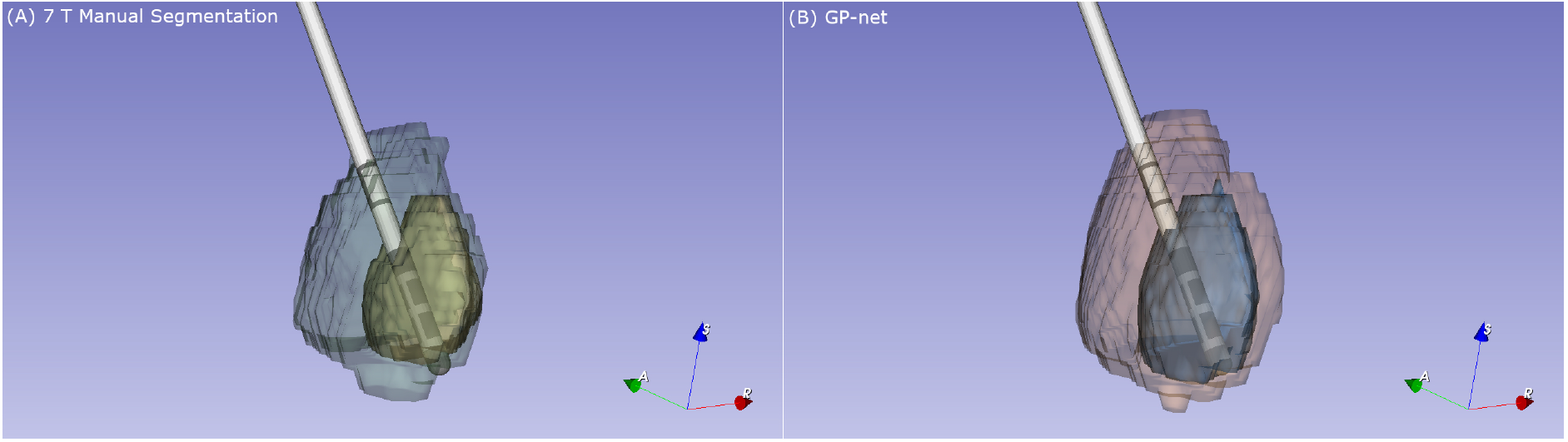
(A) 3D reconstruction of a DBS electrode placement with respect to the manually delineated GPe (green)/GPi (yellow) for a specific PD patient (age: 54). (B) 3D reconstruction of the same DBS electrode with respect to GP-net’s segmentation of the GPe (orange)/GPi (blue).

We finish this section by presenting an example which clearly demonstrates the efficacy of patient-specific segmentation over atlas-based segmentation. A subject with irregular blood vessel bifurcations which traverse into the lower GP region is presented. This subject was not part of the previous statistical analysis and was not part of the training set. Figure 8 demonstrates the manual segmentation (upper row), GP-net segmentation (middle row), and a selected atlas, Ewert 2017 (Ewert et al., 2018) (bottom row). Traversing blood vessels were manually segmented (shown in red), while the manual segmentation and automatic segmentations follow the previously used color convention. As the MNI template does not contain the irregular blood vessels, the atlas-based segmentation is stretched above the vessels in the registration process and misses the true GP region. This can further be observed in the corresponding 3D reconstructions, where the manual and GP-net segmentations near the blood vessels are correct, while the GP atlas-based segmentation is posed above them with a wide gap between, missing sections of the GPi and GPe. Table 5 summarizes the different metrics for this example.

**Figure 8.**
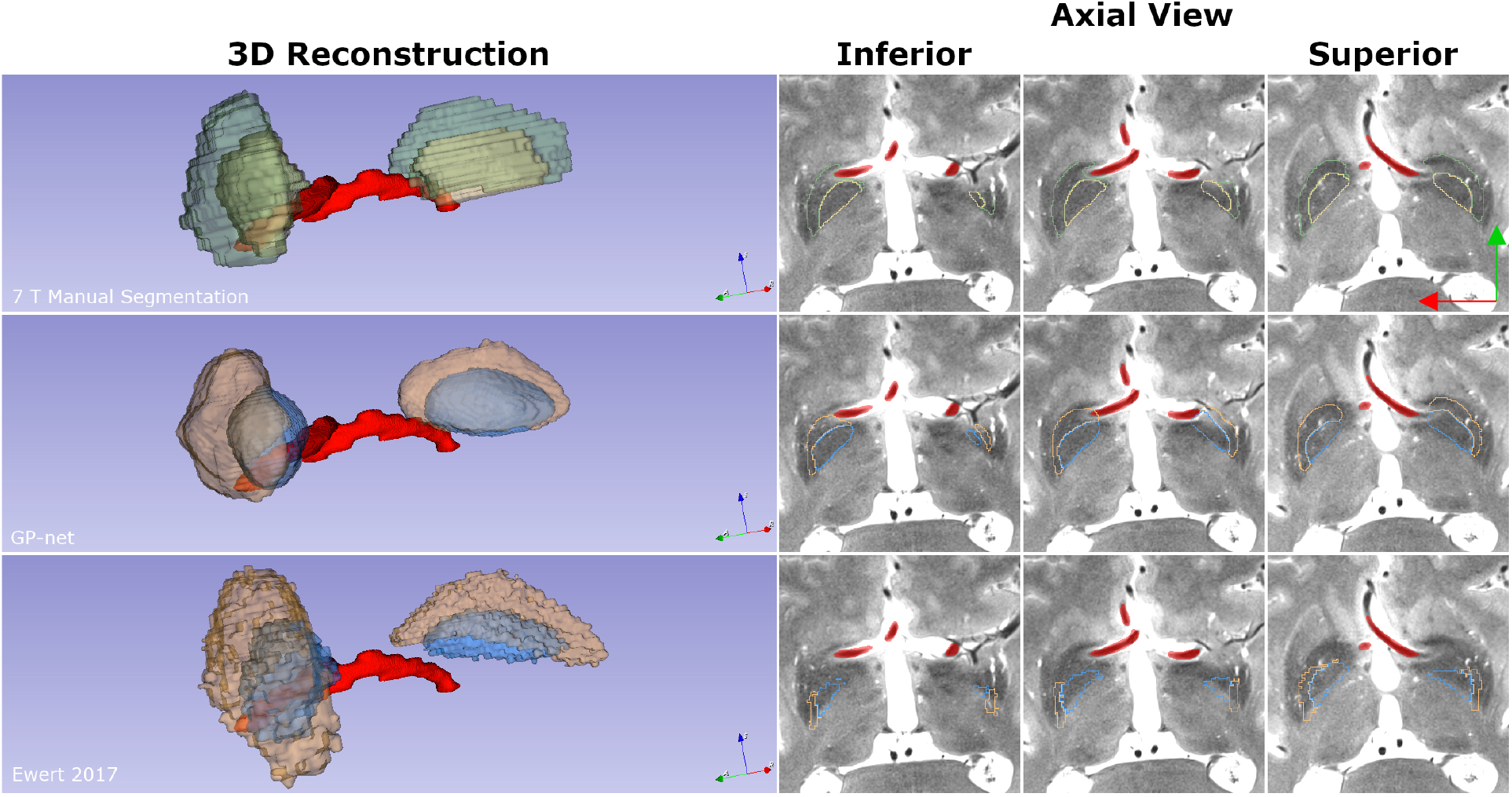
GPe/GPi segmentation of a PD patient with irregular blood vessels. Upper row, left to right: 3D reconstruction of the manual segmentation of the GPe (green) and GPi (yellow). Blood vessels are in red (smoothed only for visualization purposes, no other smoothing was applied to any structure). Different panels correspond to selected axial T2 slices, going from the inferior side to the superior side of the brain. Middle row: GP-net reconstruction (GPe in orange and GPi in blue). Bottom row: Segmentation based on the DISTAL atlas (Ewert 2017 (Ewert et al., 2018)). Other atlases produced similar results to the DISTAL atlas, and were thus omitted for brevity. Blue arrow points in the superior direction, green arrow points in the anterior direction, and red arrow points in the right direction.

**Table 5.** Measured metrics (GPe/GPi) for the example in Fig. 8. Each value is an average for both the left and right segments. Manually measured volume estimates are 1.07/0.63 [mm^3^] for the GPe and GPi, respectively.

|  | *Dice* | *MSD [mm]* | *CoM [mm]* | *Vol [mm<sup>3</sup>]* |
| --- | --- | --- | --- | --- |
| *GP-net* | 0.61/0.67 | 0.83/0.77 | 0.82/1.4 | 0.86/0.4 |
| *Ewert 2017 (DISTAL)* | 0.41/0.44 | 1.27/1.31 | 4.06/3.34 | 0.79/0.4 |

## 4. Discussion

Alterations in neuronal activity in the internal segment of the GP has been shown to be correlated with motor symptoms of PD. For example, animal models of PD have shown a characteristic increase in neuronal activities in both the STN as well as in the GPi (Obeso et al., 2001; Wichmann et al., 1994). Lesions applied to these regions have shown striking improvement in motor function. Moreover, the creation of lesions in the GPi of PD patients have been reported to improve contralateral dyskinesia and provide moderate antiparkinsonian benefits (Baron et al., 1996, 2000; Obeso et al., 2001; Vitek et al., 2003). However, adverse effects which may be caused by lesions cannot be averted once surgery is performed (Kringelbach et al., 2007; Munro-Davies et al., 1999). On the other hand, DBS was shown to provide similar clinical benefit compared to a lesion-based therapy while avoiding permanent brain damage which might occur with lesioning (Benabid et al., 1987; Obeso et al., 2001; Vitek et al., 2020). The success of DBS surgery directly relates to the accurate identification of target regions (Paek et al., 2013; Patel et al., 2015; Patriat et al., 2018; Richardson et al., 2009; Rolston et al., 2016; Welter et al., 2014), thus greatly motivating the need for an accurate, robust and reliable identification of these brain areas in an automated manner.

GP-net is a deep-learning based segmentation technique specifically tailored to an accurate and robust segmentation of both the GPe and GPi. Although GP-net was described here in the context of DBS surgery, all clinical procedures which require pre-surgery GP trajectory planning, such as DBS surgery and magnetic resonance guided focused ultrasound (Ebani et al., 2020; Miller et al., 2020; Zaaroor et al., 2018), can benefit from this method. In this work we have utilized recent advances in ultra-high MRI scanner and acquisition protocols and train the network in an end-to-end manner on pairs of 7 T T2 acquisitions and manual delineations produced by domain experts. GP-net is based on several key components: it is a U-net structure which relies on 3D convolutions (Goodfellow et al., 2016) as well as the recently introduced 3D deformable convolutions. It also relies on data augmentation, to effectively increase the size of the training set. By mirroring each T2 scan around the anterior-posterior axis, e.g. left to right (as well as the corresponding manual delineation), we effectively double the amount of training data. The trained network is able to produce fast (a few seconds per subject), accurate and reliable GPe and GPi segmentations from new 7 T T2 images.

The results presented in this work imply two key observations. First, for all the metrics considered in this work, deeplearning based GP-net was found to be superior to all the atlas-based segmentations tested. GP-net has exhibited improved average dice (scores above 0.8) indicating its ability to more accuratly capture the shape of the structure. A higher mean value along with reduced variance indicates that not only does GP-net perform better on average, it is also more stable and has far fewer outlier segmentations. This conclusion is also supported by the p-value matrices below panel (A) of Fig. 2. The first row clearly indicates that GP-net is statistically different than the atlas-based segmentations. On the other hand, the atlas-based segmentations do not significantly differ from one enother (e.g. Ewert (Ewert et al., 2018) and Pauli (Pauli et al., 2018)), which is indicative of similar performance.

Moreover, the average CoM difference, an indication of how well the structure can be localized in the brain, was measured to be on the order of ~0.7mm, which corresponds to a difference of less than two voxels on the resampled grid (0.39 mm). This CoM localization error (with respect to the manual delineations) is below the actual slice thickness of the acquired 7 T T2 volumes (1 mm). The average mean surface distance for both GPe and GPi was measured to be ~0.4 mm, which is on the order of a single voxel on the resampled T2 grid. These numerical results present a significant improvement compared to the atlas-based registrations. All atlas-based registrations achieve an average dice score of about 0.45 – 0.66 and an average CoM error larger than 1.6 mm which corresponds to 4 pixels on the resampled grid. The average MSD for the atlas-based approaches is between 0.74 – 1.32 mm, higher than the MSD reported for GP-net.

In many cases, there exists a trade-off between the precison and recall rates, as exemplified in Fig. 4, the precision rates of the atlas-based segmentation of Xiao 2017 (Xiao et al., 2017) is notably higher than its recall rates. However, such a trade-off does not seem to exist for GP-net. The rates of the other atlas-based segmentations do not exhibit such a trade-off, however these rates values are clearly lower than those of GP-net. GP-net exhibits both the highest precision and recall rates, both approaching the maximum value of 1, which further validates the superior performance of GP-net over the atlas-based segmentations.

Variability between patients has been previously reported, e.g. in (Duchin et al., 2018; Kim et al., 2019; Lenglet et al., 2012; Patriat et al., 2018). This intrinsic variability between different subjects motivates the need for patient-specific care, and suitably tailored algorthims to address this need. Standard atlases, which are typically defined in a normalized space, have shown great importance in retrospective population studies (Horn, Kühn, et al., 2017; Horn, Neumann, et al., 2017; Horn, Reich, et al., 2017; Kim et al., 2019). However, atlas-based registrations, which are ultimately based on a single depiction of the target structure (even though this depiction can be based on multiple inputs from multiple patients and modalities), cannot fully capture the intra and inter-patient variability. Intra and inter-patient variablity, as well as the need for patient-specific imaging tools is clearly exemplified by panel (D) of Fig. 2. It is sufficient to consider the volume estimations for both GPe and GPi, obtained from the manual delineations, to see that profound variability exists between patients. The ratios between the smallest and the largest meaured volume estimate correspond to an increase of 200% and 284% for the GPe and GPi, respectively. From a statistical point of view, all volume estimates of the atlas-based approaches have a consistently smaller distribution. Only GP-net exhibits a distribution which is similar to the manually measured volume distribution, for both parts of the GP. This dramatic variability in GPe/GPi volume estimates further motivates the need for a stable and consistent patient-specific, accurate segmentation tool, especially for any invasive procedure such as DBS surgery.

GP-net is trained on healthy subjects as well as PD and ET patients and with a large age distribution. This training process allows the network to encounter and learn from a large variety of different pathologies (Fig. 3A-B) and across varying age groups (Fig. 3C-D), thus allowing stable and robust segmentation performace. Robust performance across different pathologies is further exemplified by the p-value matrices below Fig. 3A-B. These conclusions are further supported by the average and standard deviation values presented in Table 4. P-value matrices below Fig. 3C-D show no statistical significance between the different age groups (calculated for GP-net). This ability of GP-net to generate accurate segmentations over different clinical pathologies and age groups suggests that GP-net might play an important role in clinical DBS targeting or similar applications for a variaty of patients populations.

One of the key advantages of the proposed method is that GP-net relies solely on 7 T T2 scans to perform its inference. No registrations are involved in its process and, therefore, operates directly in the patient’s system coordinates. Atlas-based segmentations on the other hand, are greatly affected by the registration process, which can adversly affect the outcomes of DBS surgery. Unfortunately, registration errors cannot be modelled easily, are typically unpredictable, and often have large variance (Kim et al., 2019). Moreover, the registration process often involves several successive registrations, which tend to accumulate and increase the registration errors with each step. Due to these factors, registrations often must be verified manually, as was done in this work for the atlas-based registrations. This dramatically prolongs the processing time per patient.

Although the atlas based registrations and final segmentations were performed on the same grid as GP-net, they are visually more pixelated, as no spatial filtering is performed on the acquired T2 volume to infere the segmentations. Contrary to this, GP-net operates directly on the 0.39 mm^3^ isotropically resampled grid of the T2 volume, and by the use of 3D convolution filters it is able to produce smooth, deliniated segmentations, and higher quality structure detection of both parts of the GP.

Since atlas-based registrations rely on a single (often average) depiction of the target structure, such a process cannot truly account for different intra-patient variabilities in a true patient-specific manner, which is clearly exemplified by the segmentation results depicted in Fig. 8. The middle row of Fig. 8 shows that GP-net is able to segment the GP, while accounting for the blood vessels, even though GP-net was not trained on such irregular cases. GP-net is able to account for this irregular vessel shape and still be able to produce clear GP segmentations which align with the underlying 7 T scan anatomy, and is relatively close to the manual segmentation. On the other hand, the atlas-based segmentation presented in the lower row of Fig. 8 is greatly affected by the registration process of the MNI template (which does not contain such vessels) and thus erroneously segments a large portion of the GP. The large CoM difference for the atlas based segmentation, presented in Table 5, further validates and quantifies this misalignment. This example indicates the true potential of deep-learning based approaches such as GP-net, in being a patient-specific automatic segmentation technique, even for irregular and unique cases, without any additional training.

As was mentioned before, the internal and external segments of the GP are separated by a thin lamina layer. Often when using MER during DBS surgery, the lamina layer is characterized by the absence of somatodendritic action potentials, which characterize both the GPe and GPi, each with its own characteristic firing pattern (Baron et al., 1996, 2000; Lozano & Hutchinson, 2002). Thus, a clear and reliable visual depiction of the lamina can also be of invaluable importance to DBS surgery. Even when using high contrast 7 T T2 scans, this border is not always clearly presented in the scans. However, the overall GPe and GPi structure can be infered, as presented in this work. A possible way to achieve lamina depiction is by accurately detecting the two GP compartments, and to indirectly infer the lamina border from both segmentations. This direction is a matter of future research and extension for GP-net.

GP-net has two potential limitations which are attributed to its deep learning-based framework. First, at this stage, GP-net was only tested on 7 T T2 data, which was acquired with the protocol described in this manuscript, as GP-net was trained solely on this type of data. This is a common limitation to many deep learning-based architectures and achieving increased robustness in the face of changing datasets (e.g. varying signal to noise ratios, resolution, ect.) is a matter of ongoing research in the machine learning community (often denoted as domain shift or domain transfer). Extending this work to operate on standard clinical images (1.5 T - 3 T) is the subject of future work. Second, deep learning frameworks currently lack interpretability. In some cases, the inference might result in sub-optimal performance (e.g. in panel (A) of Fig. 2, two GPe dice scores for GP-net are below 0.6). In these cases, it is not always easy to understand why the network reacted the way it did. Fortunately, as was statistically veryfied in this work, such occurrences seem to be rare. Exploiting standard CNN visualization tools can potentially educate the user on the internal workings of GP-net.

The clinical potential of GP-net is clearly exemplified in Fig. 7, which presents an excellent match between the implanted DBS electrode and the manual versus GP-net’s segmentations of both GPe/GPi, for a representitive PD patient. GP-net will provide the clinical team the ability to have a much better understanding of the correlation between lead locations and outcomes. For DBS surgery, both the center of mass of the DBS target, as well as the accurate identification of its borders are of great importance. This figure, supported by the reported results for CoM distance and MSD, show that an accurate depiction of the DBS target clearly achieves this goal. This accurate identification of both compartments of the GP can lead to a reduction in the number of MER penetrations and, subsequently, reduced time in the operating room. One barrier as to why patients chose not to proceed with DBS surgery is the necessity to be awake during the MER portion of the surgery (Falconer et al., 2016). Reducing time needed for MER would have a profound effect on increasing the number of patients willing to proceed with surgery. Additionally, reducing the number of microelectrode penetrations decreases the probability of intraoperative and/or postoperative complications (Ben-Haim et al., 2009; Binder et al., 2005). Accurate identification can also contribute to reliable and accurate DBS lead placement, which has been associated with improved clinical outcomes in leads placed in the STN (Richardson et al., 2009). This often leads to a reduced amount of programming time in the clinic necessary for optimizing symptom reduction and an overall improvement in quality of life for the patient (Hell et al., 2019).

## 5. Conclusions and future work

In this work we presented GP-net, a deep-learning based neural network for the segmentation of both the GPe and GPi. GP-net is able to produce accurate and reliable segmentations in a fully automated manner from 7 T T2 MR acquisitions, both for healthy subjects as well as PD and ET patients and with a large age distribution. The network is trained end-to-end on pairs of acquired 7 T T2 scans and corresponding manual delineations of both GPe and GPi. We have shown, both qualitatively and quantitatively, that GP-net outperforms state-of-the-art atlas-based segmentations and produces stable and consistent high quality patient-specific segmentations, while reducing potential biases.

GP-net is tailored for the segmentation of the GP (both internal and external segments). However, it can be extended to segment additional subcortical structures which are of interest for DBS surgey, such as the STN, Red Nucleus, and Substantia Nigra. This extension is currently being investigated in our group.

With the incorporation of advanced DBS pre-surgery targeting and post-surgery lead localization tools and software, 7 T MRI based approches, either for training or for deployment, have great potential in becoming clinical standards, especially now that the 7 T MRI is FDA approved for standard clinical applications. In this scenario, the use of fully automated segmentation software may prove to be very advantageous, leading to an accurate, fast, easy and reliable visualization tool, contributing to an improved surgical procedure and patient experience.

## Availability of data

The data that support the findings of this study are available on request from the corresponding author. The data are not publicly available due to privacy or ethical restrictions.

## Acknowledgements

This work was funded by the following national institution of health grants: R01 NS081118, R01 NS113746, P50 NS098753, P30 NS076408. Additional support by NSF (GS) and Department of Defense (GS) is also aknowledged.

## Footnotes

i Intraoperative microelectrode recording (MER) could be considered as well. MER for the GP is significantly less understood than for the STN for example, is time consuming, and requires expertise.

ii Code will be publicly available upon publication

